# Milk losses linked to udder health treatments at dairy farms with automatic milking systems

**DOI:** 10.1101/2021.01.04.425263

**Authors:** Ines Adriaens, Igor Van Den Brulle, Ben Aernouts

## Abstract

Milk losses caused by intramammary infections (IMI) have a massive impact on farm profitability and sustainability. In this study, we analyzed milk losses from 4 553 treated IMI cases at 41 AMS dairy farms. Milk losses were estimated based on the difference between the expected and the true production. To estimate the unperturbed lactation curve, we applied an iterative procedure using the Wood model and a variance-dependent threshold on the milk yield residuals. We calculated milk losses both in a fixed window around the first treatment day of each IMI case and in the perturbations corresponding to this day, at cow level as well as at quarter level. In a fixed time window of day -5 to 30 around the first treatment, the absolute median milk losses per case were 101.5 kg, highly dependent on the parity and the lactation stage with absolute milk losses highest in multiparous cows and at peak lactation. Relative milk losses expressed in percentage were highest on the treatment day, and full recovery was often not reached within 30 days from treatment onset. In 62% of the cases, we found a perturbation in milk yield at cow level at the time of treatment. On average, perturbations started 8.7 days before the first treatment and median absolute milk losses increased to 128 kg milk per perturbation. Mastitis is not expected to have equal effects on the four quarters so this study additionally investigated losses in the individual udder quarters. We used a data-based method leveraging milk yield and electrical conductivity to project the presumably infected quarter and compared losses with the average of presumably non-infected quarters. Median absolute losses in a fixed 36-day window around treatment varied between 50.2 kg for front and 59.3 kg for hind infected quarters compared to respectively 24.7 and 26.3 kg for the median losses in the non-infected quarters. Also here, these losses depended on lactation stage and parity. Expressed proportionally to expected yield, the relative median milk losses in infected quarters on the treatment day were 20% higher in infected quarters with a higher variability and slower recovery. In 86% of the treated IMI cases, at least one perturbation was found at the quarter level. This analysis confirms the high impact of IMI on milk production, and the large variation between quarter losses illustrates the potential of quarter analysis for on-farm monitoring at farms with an automated milking system.

**Highlights:**

- Milk losses were estimated for treated cases of intramammary infections
- Milk losses were highly variable across cases with a median of 101 kg
- We found large differences between infected and non-infected quarters
- Quantification of milk losses can be the basis for better udder health management

## INTRODUCTION

Intramammary infections (**IMI**) lead to poor profitability and decreased dairy farming sustainability, having significant impact on animal welfare, efficiency of production and milk quality. Since problems are often detected later in farms with automated milking systems (**AMS**) compared to conventionally milking operations, the impact of IMI is even higher at AMS farms (Barkema et al., 2015; Deng et al., 2020).

Quantification of milk losses and milk yield dynamics raises the farmers’ awareness of the impact of IMI on a factual basis. This lays the groundwork for improved detection and treatment decisions, as such limiting their negative impact. In the past, most studies either used low frequency data to study these milk losses, or relied on experimental infections limiting the number and representativeness of IMI cases included (Gröhn et al., 2004; Bar et al., 2008). A solution is to analyze the impact of IMI based on commercial farms’ treatment registers and milk meter data.

Unfortunately, high-quality datasets comprising both treatment records and high-frequency on-farm milk meter data are scarce. Reliability of treatment registers, moreover, is often hampered due to poor registration by farmers, both in quality and quantity (Stevens et al., 2016), while only in the past decennia high-frequent milk production data became more frequently available on farm. The latter has tremendously improved with the increased use of AMS that record and store milk yield per milking as well as per quarter in an automated way.

With high-frequency milk production time series available, it becomes possible to accurately estimate milk losses associated with health problems. To this end, a good estimation of the expected milk production in absence of perturbations is needed. For example, we (Adriaens et al., 2018, 2020, 2021) and others (Poppe et al., 2020; Ben Abdelkrim et al., 2021) developed methodologies to predict milk yield in an unperturbed state. These predicted trajectories and their corresponding deviations can subsequently be used to study resilience, for improved monitoring and precision phenotyping as well as for the characterization of milk yield dynamics and losses during challenges. In the case these challenges involve udder health problems, the milk yield of the 4 individual udder quarters can be mutually compared next to studying the cow-level milk yield. Quarter separation additionally provides a unique opportunity to study losses and dynamics in the different quarters separately and to distinguish local (i.e., related to the local inflammatory reaction and toxins) and systemic (i.e., related to the general sickness) effects of the infection (Adriaens et al., 2018).

A better quantification of the actual milk losses during IMI provides the necessary knowledge to further raise awareness among farmers and veterinarians on the impact of udder health on farm, and by extension, on dairy production’s sustainability through improved profitability and decrease antimicrobial use. Distinguishing between the cow and quarter level additionally leverages new opportunities for better use of available on-farm data for monitoring and decision support. Starting from an elaborate data set that includes both the high-frequency sensor data and treatment registers from modern AMS dairy farms, this study aimed to describe milk losses related to treatments of IMI. To this end, we studied cow and quarter level milk yields and losses both in a fixed period around the first treatment and in perturbations associated with these treatments.

## MATERIALS AND METHODS

### Data collection

This study relied on two main sources of data: high-frequency sensor data and udder health treatment registers. For the sensor data, back-up files of the AMS software were collected on 50 modern dairy farms with a Lely (N=20, Maassluis, The Netherlands) or DeLaval (N=30, Tumba, Sweden) AMS in Belgium and The Netherlands during a farm visit. For DeLaval farms, additional back-ups were downloaded at intervals of 180 to 400 days depending on the AMS software settings. This procedure ensured access to minimal 1 year of uninterrupted longitudinal sensor time series for the analysis. During the farm visits, we also copied the treatment registers related to udder health problems. These treatment registers were either handwritten on paper, recorded in a spreadsheet, included in the AMS software or entered in separate farm management software (Unifarm, Uniform-Agri, Assen, The Netherlands).

### Editing and selection of sensor data

Raw data tables were exported as flat files from the AMS software back-up files using T-SQL and Microsoft SQL Management Studio (Microsoft, Redmond, USA). The different tables were horizontally (combining cow identification (**ID**), dates, etc. with the sensor records) and vertically (combining different back-up files of the same farm) merged and further processed using Matlab 2019a (MathWorks, Natick, MA, USA). Per farm, two data sets of sensor data were compiled: one set containing the daily milk production records at cow level (**CMY**), and a second set containing the sensor data per individual milking at quarter level, including the milk yield (**QMY**) and electrical conductivity. The first data set of CMY contained records from the installation date of the AMS system onwards for all farms. The daily data coming from DeLaval AMS represent the sum of the milk yields produced each calendar day (i.e. 24 hours), while the Lely AMS software, before storage, corrects for the varying number of milkings per day per cow when milking with AMS. To obtain a similar variance as for the Lely milk production data, the daily milk production time series of DeLaval farms were smoothed with a moving average using a span of 3 days before the analysis (Adriaens et al., 2020). The second data set for Lely AMS systems contained all data since the installation of the AMS and thus matches with the daily data, whereas for DeLaval systems the second data set typically comprised a shorter period compared to the daily data. This is because individual milking information in the DeLaval software is only stored for a limited amount of time, typically 180 or 400 days depending on the software settings. For each farm and each data set, we kept only the lactations with enough data to reliably estimate milk losses. To this end, we selected lactations that started within 5 days after calving (days in milk, **DIM**) and for which at least 150 days of milk production data were available. Lactations for which data were missing during more than five successive days were also excluded from the analysis. Only the first 305 days of milk production were considered to avoid the lactation curve being affected by pregnancy or impending dry-off.

### Expected milk production

To calculate milk losses as the expected versus the actual milk production, first a reliable estimation of the expected production must be obtained. For the CMY analysis, the method described in Adriaens et al. (2021) was used. This is based on an iterative fitting of the Wood model (Wood, 1967) in which records that are part of a perturbation are removed to obtain a fit unaffected by perturbations. For the QMY data, the methodology for estimating the unperturbed lactation trajectory was adjusted to deal with the higher variability in production and to account for the varying milking intervals. To this end, we evaluate the milk yield residual against a threshold of 1.6 times the standard deviation of these residuals as described below. For the QMY, following steps were implemented, visualized in Figure 1:

1. Upper panel Figure 1. The QMY was transformed to QMY_t_, expressed in kg milk per 24 hours by dividing the QMY by the milking interval and multiplying it by 24 hours (pink circles). The multiplication aids in stabilizing the parameters of the Wood model in the iterative fitting procedure, while the correction for milking interval ensures that the variability caused by different intervals between milkings does not bias the lactation curve shape.
2. To correct for outliers with very high milk yields e.g. caused by a failure of the previous milking, the QMY_t_ was smoothed with a median smoother (dark grey solid line) on a window of 18 measurements (+/- 2 weeks at a milking frequency of 2.6 milkings per day). The QMY_t_ above 4 times the median average distance (MAD) between the QMY and this median smoothed curve (light grey solid line and green squares) were deleted for the estimation of the model in de next steps.
3. An initial Wood curve (QMY_t_ = a * DIMb * e^-c*DIM^) is fitted on all the remaining QMY_t_ of the lactation to obtain the initial unperturbed lactation curve ULC1. The residuals from this initial model are calculated as QMY-ULC_1_, from which the standard deviation (SD_1_) and the root mean squared error 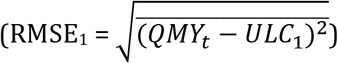 were computed.
4. QMY_t_ below 1.6 * SD_1_ were deleted (orange squares) except when they occurred in the first 5 days of lactation to ensure correct fitting of the non-linear shape in the beginning of the lactation curve. Accordingly, a new non-linear Wood curve (ULCi, orange dashed line) was fitted on the remaining QMY_t_. Again, the QMY_t_ for which the residuals were below 1.6 * SD_i_, but which were not in the first 5 days of lactation were deleted. The RMSE_i_ of the remaining residuals was computed.
5. This refitting and deletion iterations were continued until the difference in RMSEi between the current and the previous iteration was less than 0.1 kg of milk.
6. The final model (blue solid line in upper panel) was retransformed, by multiplying it with the milking interval and dividing it by 24, to obtain the expected milk production for that udder quarter for each milking event (unperturbed lactation curve, ULCF, blue solid line in middle panel Figure 1).

**Figure 1.**
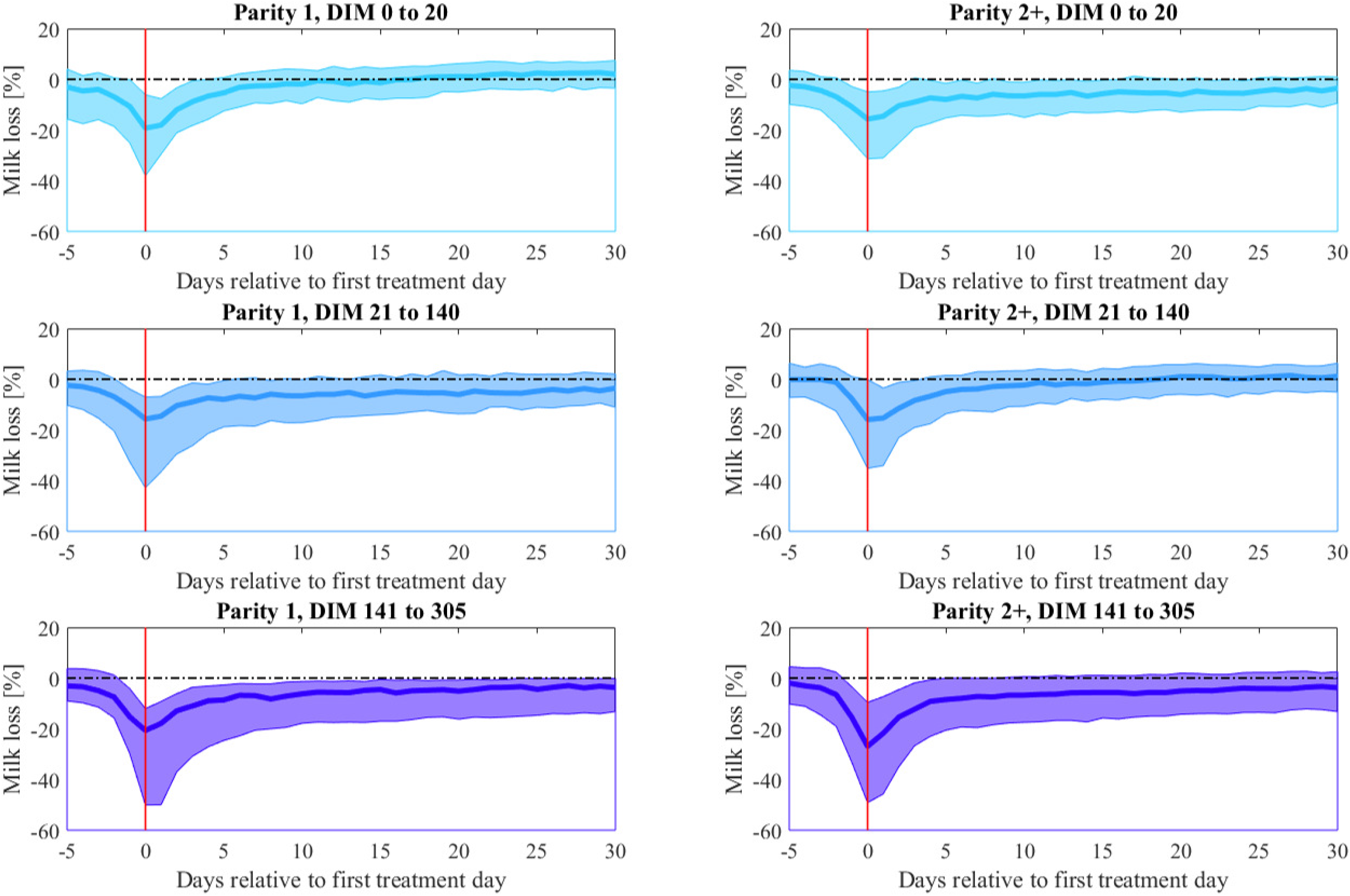
Visualization of the different steps to obtain the unperturbed milk production curves at quarter level. The upper panel shows the transformed quarter milk yields (QMY_t_), the data excluded because of outlying values above 4 times the median average distance (MAD) from a 18 measurement median window smoothing (grey solid lines, green squares), the measurements excluded because they were below the threshold in estimating the unperturbed curves (orange dots), and the final model ULCF (blue solid line). The middle panel shows the raw QMY data and retransformed ULCF. The lower panel shows the relative residuals (% per day) from this model, the median smoothed relative residuals and the 80% threshold (red solid line) applied to identify perturbations.

Milk losses are calculated as the difference between the actual production and the expected production. Dividing the absolute milk losses (i.e. kg of milk) by the expected milk yield renders the relative milk losses (% milk loss, lower panel in Figure 1).

### Identification of perturbations in milk yield

For this study, we calculated milk losses at cow and quarter level both in a fixed window around the day of the first udder health treatment (treatment day, **TD**), and in perturbations in which the TD was included. The latter provides additional information on when the deviation in milk yield starts and on its true duration. Perturbations are considered as periods in which the milk production is consistently below the expected milk production. For the CMY data, again similar criteria as in Adriaens et al. (2020, 2021) were applied. In brief, at cow level, a perturbation was defined as an episode for which the CMY was below the expected yield (ULCF) for at least 5 consecutive days, with at least one day below 80% of the expected production. These criteria are the result of discussions with veterinary experts and are motivated in Adriaens et al. (2020, 2021). For the QMY, the higher variability compared to CMY data needed to be considered. Therefore, perturbations were identified on the 7-day median smoothed relative residuals (Figure 1, lower panel, orange line) rather than the raw data. A perturbation was considered as a period in which (1) the median smoothed residuals were below 0 for at least 7 consecutive days, and (2) for which at least 5 milkings were below 80% of the expected yield (Figure 1, lower panel, red line). Perturbations between individual udder quarters of the same cow were compared and an overlap was identified when the minima of the perturbations were separated in time by less than 50% of the average length of those perturbations.

### Editing of treatment registers and selection of IMI cases

In a first step, all available treatment registers were digitalized and their format standardized to obtain a well-organized list containing at least the animal ID and date for each treatment. Records with no clear ID, no date or that were not related to IMI were deleted. Only the first treatment of the IMI was considered, and records from the subsequent days were removed. If there was a gap of more than 10 days between 2 treatments, they were considered as different treated IMI cases, further referred to as IMI cases. Next, the cow IDs of the treatment registers were coupled to the unique cow IDs in the milk yield data sets. Only IMI cases with a TD that occurred within the time span of the available milk yield data and before 305 DIM after calving were kept, because only for these IMI cases the milk losses could be calculated. As the timespan for the CMY data was longer than for the QMY data, the IMI cases included for the analysis of milk losses at quarter level were a subset of those included for the cow-level analysis. Based on the quarter level data (QMY and electrical conductivity), we identified the presumably infected quarter (**IQ**) to allow comparison of milk losses between the IQ and the presumably non-infected (**NIQ**) quarters. To distinguish the IQ from the NIQ, the relative milk losses expressed in % in each quarter were calculated over a period of 36 days (day −5 to 30 from the TD). Next, following criteria were applied: 1) the IQ is the quarter for which the relative milk losses, expressed in percentage, in the 36-day period was within 1% difference from the minimum of relative milk losses of the 4 quarters, and 2) the inter quarter ratio of that quarter’s electrical conductivity was higher than 1. This inter quarter ratio per quarter was calculated as the average electrical conductivity of that quarter divided by the average of the other 3 quarters in the first 10 days after the TD. As the amount of IMI cases recorded and thus, the reliability of the treatment registers was too variable across the different farms, the data of each farm were not analyzed separately, neither was it possible to make statements on IMI incidence rates or the milk losses at farm level.

### Milk loss analyses

Milk losses were calculated both in a fixed window of 36 days around the TD (day −5 to day 30) and in perturbations that included the TD. We considered both milk losses expressed in absolute numbers (kg milk lost = produced – expected ULCF), further referred to ‘absolute milk losses’ and in relative numbers (% milk loss, ‘relative milk losses’). The calculation of the milk losses in a certain period (fixed window or perturbation) for the QMY data was based on the raw data to account for the true milk yield, and not on the median smoothed data used for detection of perturbations. Because the distribution of the milk losses did not have a Gaussian shape, we report their median, first quartile and last quartile. The milk production dynamics are known to differ across parities and lactation stages (Wilson et al., 2004; Hertl et al., 2014; Heikkilä et al., 2018). Therefore, we report the milk losses in 3 lactation stage classes (**LS1**: DIM 0 to 20, **LS2**: DIM 21 to 140 and **LS3**: DIM 141 to 305) and 2 parity classes (**P1**: first parity vs. **P2**: higher parities). For the analyses at quarter level, we distinguished losses in IQ and the average of the NIQ, and we used a paired t-test to compare their level at the different moments relative to the TD. For the perturbation analysis at quarter level, we considered the milk losses for the duration of the perturbation detected in the IQ for all 4 quarters.

## RESULTS

### Data overview

From the 6 299 IMI cases that were identified based on the treatment registers and for which animal IDs were available, 4 553 IMI cases overlapped with the CMY data. Only for these we could calculate the milk losses at cow level. These 4 553 IMI cases originated from 2 387 unique cows, 2 999 lactations and 41 farms (25 Delaval, 16 Lely). This means that for the animals that had at least one IMI treatment registered, on average 1.9 IMI cases were recorded, corresponding to 1.5 IMI cases per lactation. The time between the first and last treatment of a farm included in the analysis was on average 3.7 years, but the variability between farms was very high, ranging from 250 days to 14 years. Further information on the number of IMI cases, cows and lactations at farm level is given in Table 1, together with their distribution over lactation stages, parities, and seasons. We had more treatments in July to September compared to the other months, and proportionally more IMI cases in early lactation. With the covered period for QMY data in DeLaval software back-ups being shorter, only 3 369 cases from 37 farms (21 DeLaval, 16 Lely), 1 882 cows and 2 296 lactations could be included for the analysis at quarter level. In this data set, we see a similar proportion of IMI cases for parity, lactation stage and season as for the cow-level data set, resulting in a higher percentage of IMI cases treated in the first 20 DIM and in the summer (July to September).

**Table 1:** Overview of the treated IMI cases that overlapped with the cow-level milk yield (CMY) and the quarter milk yield (QMY) data per farm and the proportion of cases per lactation stage (LS), parity (P) and season over the entire data set.

|  | Treated IMI cases |  |
| --- | --- | --- |
|  | CMY | QMY |
| **No. of IMI cases** (mean ± SD, [range]) | 111 ± 191, [1;1066] | 91 ± 147, [3;790] |
| **No. of unique cows** (mean ± SD, [range]) | 58 ± 78, [1;431] | 50 ± 65, [3;355] |
| **No. of unique lactations** (mean ± SD, [range]) | 73 ± 108, [1;582] | 62 ± 488, [3;467] |
| **Proportion of IMI cases per LS<sup>1</sup>** (% per day) |  |  |
| - LS1 | 0.66 | 0.61 |
| - LS2 | 0.37 | 0.38 |
| - LS3 | 0.25 | 0.26 |
| **Proportion of IMI cases per parity** (%) |  |  |
| - P1 | 19.3 | 15.7 |
| - P2 | 80.7 | 84.3 |
| **Proportion of IMI cases per season** (%) |  |  |
| - January to March | 22.5 | 22.3 |
| - April to June | 23.1 | 23.7 |
| - July to September | 30.1 | 30.5 |
| - October to December | 24.3 | 23.5 |

<sup>1</sup> LS: lactation stage, LS1 = days in milk (DIM) 0 to 20, LS2 = DIM 21 to 140, LS3 = DIM 141 to 305; P: parity, P1 = parity 1, P2 = parity 2 and higher

Table 2 gives an overview of the milk production characteristics of the included farms, calculated from all lactations for which data of at least the first 150 DIM were available. Because the period for which data were available varied across farms (e.g. for some farms only 1 or 2 years of data could be included, limiting the amount of selected lactations), this is not an exact representation of the farm characteristics, but it rather represents the data composition at farm level used for the analysis.

**Table 2.** Overview of the farm characteristics for the cow-level (CMY) and quarter milk yield (QMY) data sets in the period covered by the treatment registers

|  | CMY | QMY |
| --- | --- | --- |
| | mean ± SD, [range] | mean ± SD, [range] |
| **No. of unique cows** | 409 ± 773, [26;5079] | 262 ± 203, [26;1036] |
| **No. of lactations** | 704 ± 935, [26;5656] | 510 ± 491, [26;2748] |
| **Covered time period** (days) | 1390 ± 1126, [255;4970] | 1023 ± 878, [203;4605] |
| **Average daily milk yield** (kg) | 32.2 ± 3.7, [22.3;43.6] | --- |
| **Average per milking yield** (kg) | --- | 11.8 ± 1.4, [8.7;14.8] |
| **Average 305-day milk yield** (kg) | 9856 ± 1147, [6784;13411] | 10105 ± 1045, [7557;13399] |
| **Milking frequency** (milkings/day) | --- | 2.8 ± 0.24, [2.2;3.3] |

### Milk losses at cow level

#### Milk losses in a fixed window around treatment

We found that on average, milk losses between TD −5 to TD +30 summed up to 164 ± 204 kg per IMI case (mean ± SD), while the median milk losses were 101 kg and the 25 and 75 percentiles were 38 and 215 kg. Table 3 gives the absolute milk losses in a fixed window of TD −5 to TD +30, whereas Figure 2 presents the relative losses per day, split per parity and lactation stage class (right, P1; left, P2; up to down LS1 to LS3). Table 3 and Figure 2 show that none of the distributions have a Gaussian shape, but they are skewed with a mode smaller than the medians and means and a heavy tail towards higher losses. For example, for the absolute milk losses in the fixed window around the TD, the mode was 69 kg, while the median and mean were respectively 101 and 164 kg milk, implying that proportionally more IMI cases had smaller losses and a small number of IMI cases had extremely high milk losses. From Table 3, it can be concluded that the absolute milk losses increase with lactation stage in first parity cows. In higher parity cows, the absolute milk losses for IMI cases treated between day 21 to 141 were higher than in the early or late lactation stages. The absolute milk losses were lower in first parity cows compared to higher parity cows. In the early and late LS, the milk losses were about equal across parities. In 76.1% of the IMI cases, there were milk losses registered in the 5 days before TD, resulting in median milk losses of 11 kg in the 5 days before TD considering all parities and lactation stages together. Only for IMI cases detected in the first 20 DIM of 2nd and higher parity cows (P2, LS1), more than 25% of the IMI cases had no losses before treatment. The milk losses in the days after TD reflect the severity and recovery rate of the IMI cases. Overall, median milk losses add up to 75.6 kg milk in the 30 days after the TD. However, mainly IMI cases in the first 20 DIM seem to give less milk losses, as no milk losses were found when considering the entire 30-day period after TD in more than 25% of the IMI cases. Also here, the absolute losses were highest for higher parity cows in the second lactation stage (DIM 21-140).

**Table 3.**
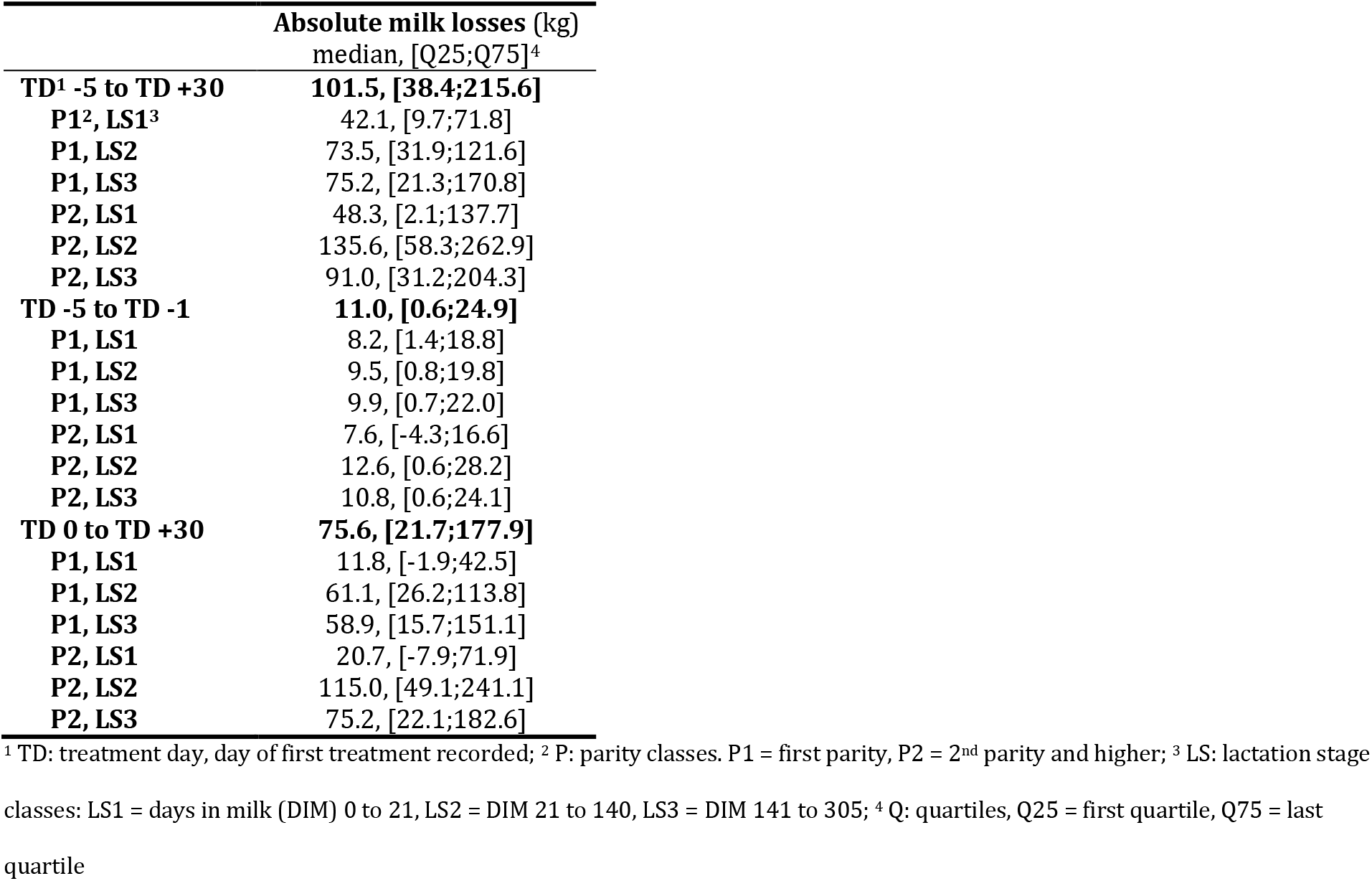
Absolute milk losses in a fixed window of 36 days around the first treatment day (TD) calculated from cow-level milk yields (CMY) in different lactation stage and parity classes. The milk losses are expressed as the amount of milk lost, and thus in positive numbers.

|  | Absolute milk losses (kg)<br>median, [Q25;Q75]<sup>4</sup> |
| --- | --- |
| **TD<sup>1</sup> -5 to TD +30** | **101.5, [38.4;215.6]** |
| P1<sup>2</sup>, LS1<sup>3</sup> | 42.1, [9.7;71.8] |
| P1, LS2 | 73.5, [31.9;121.6] |
| P1, LS3 | 75.2, [21.3;170.8] |
| P2, LS1 | 48.3, [2.1;137.7] |
| P2, LS2 | 135.6, [58.3;262.9] |
| P2, LS3 | 91.0, [31.2;204.3] |
| **TD -5 to TD -1** | **11.0, [0.6;24.9]** |
| P1, LS1 | 8.2, [1.4;18.8] |
| P1, LS2 | 9.5, [0.8;19.8] |
| P1, LS3 | 9.9, [0.7;22.0] |
| P2, LS1 | 7.6, [-4.3;16.6] |
| P2, LS2 | 12.6, [0.6;28.2] |
| P2, LS3 | 10.8, [0.6;24.1] |
| **TD 0 to TD +30** | **75.6, [21.7;177.9]** |
| P1, LS1 | 11.8, [-1.9;42.5] |
| P1, LS2 | 61.1, [26.2;113.8] |
| P1, LS3 | 58.9, [15.7;151.1] |
| P2, LS1 | 20.7, [-7.9;71.9] |
| P2, LS2 | 115.0, [49.1;241.1] |
| P2, LS3 | 75.2, [22.1;182.6] |

<sup>1</sup> TD: treatment day, day of first treatment recorded; <sup>2</sup> P: parity classes. P1 = first parity, P2 = 2<sup>nd</sup> parity and higher; <sup>3</sup> LS: lactation stage classes: LS1 = days in milk (DIM) 0 to 21, LS2 = DIM 21 to 140, LS3 = DIM 141 to 305; <sup>4</sup> Q: quartiles, Q25 = first quartile, Q75 = last quartile

**Figure 2.**
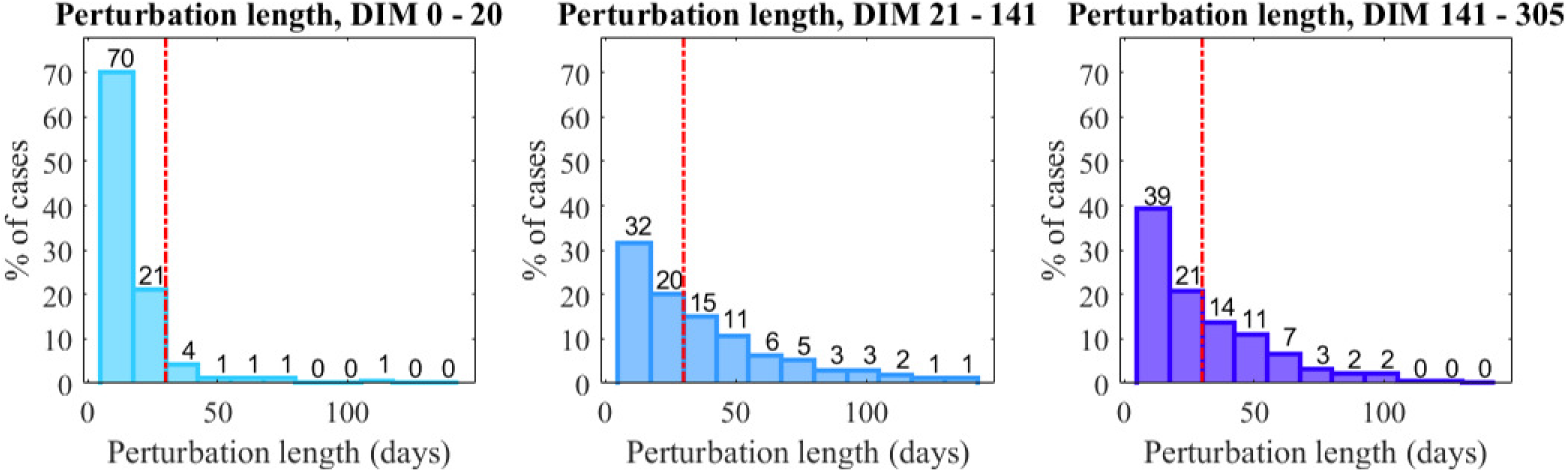
Median, 25 and 75 percentiles of the relative milk yield losses expressed in % milk losses over time in a fixed window of 36 days around the first treatment (red vertical line, day 0 of treatment) in different lactation stage and parity categories. The milk losses are shown in negative figures representing the true direction of the change during intramammary infections.

Figure 2 shows that in all lactation stage and parity classes a clear perturbation is present around the TD. Relative milk losses increase with lactation stage for both parity classes and in most classes the median relative losses do not reach 0 within 30 days after the TD. However, for first parity IMI cases before DIM 20 and for IMI cases in higher parities around peak production (LS2), milk yield seems to recover more rapidly, with the median relative milk losses reaching 0% at day 11 and day 16 respectively, demonstrating a shorter impact of IMI during these IMI cases. These figures also illustrate the variability in and skewed distributions of milk loss dynamics, with the spread at day 0 largest in all P/LS classes.

### Milk losses in perturbations

Because of the high variability in milk losses across IMI cases, we also analyzed the true impact of IMI on CMY based on the milk yield perturbations including the TD. As we set criteria for what was considered a perturbation, this part of the analysis is biased towards more severe IMI cases, but it considers the true duration of the IMI cases’ impact. A perturbation was found in 2 823 out of the 4 553 cases (62.0%), which means that for 38.0% of treatments CMY did not drop below 80% of the expected yield for at least one day, or the perturbation was shorter than 5 days. The length of the perturbations (Figure 3) varied between 5 and 190 days. Overall, shorter perturbations were more often identified in early lactation compared to later lactation stages. Over all parities, 91, 52 and 60% of the perturbations were 30 days or shorter (red vertical line in Figure 3) when the TD was in LS 1, 2 or 3 respectively. The perturbations started on average 8.7 ± 12.9 days before treatment, with the mode (most frequent value, 15.1%) being 1 day before treatment, demonstrating again the highly variable effect of IMI on milk yield.

**Figure 3.**
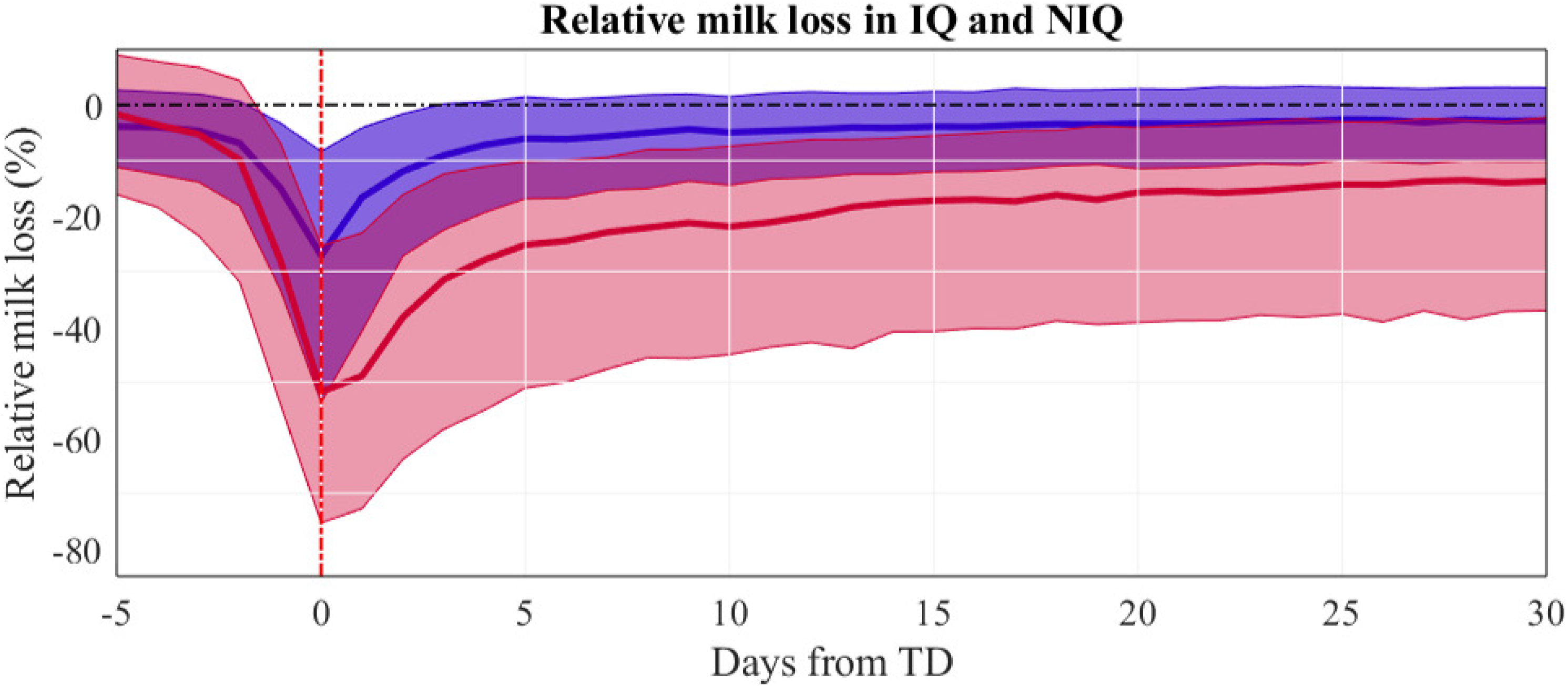
Frequencies of perturbation length around treatments for intramammary infections (IMI) in different lactation stages. The red dashed line shows the perturbations above and below 30 days of length.

The median absolute milk losses in the perturbations including a TD summed up to 128 kg, which is higher than when using a fixed window (101 kg). When comparing milk losses across lactation stages and parities, the same trends as for the milk losses in the fixed window around TD were observed (Table 4). Absolute milk losses in the first parity were lower than in higher parities.

**Table 4.** Absolute and relative milk losses in perturbations at cow level corresponding to cases of intramammary infections. The milk losses are expressed as the amount of milk lost, and thus in positive numbers.

|  | Absolute milk losses (kg)<br>median, [Q25;Q75]<sup>3</sup> | Relative losses (%)<br>median, [Q25;Q75] |
| --- | --- | --- |
| **Overall** | **128.0, [59.2;296.5]** | **17.2, [12.9;23.2]** |
| **P1<sup>1</sup>, LS1<sup>2</sup>** | 37.9, [26.2;69.6] | 16.2, [12.6;22.5] |
| **P1, LS2** | 110.9, [58.6;211.9] | 15.4, [11.9;19.8] |
| **P1, LS3** | 118.8, [59.5;237.0] | 17.2, [13.0;23.5] |
| **P2, LS1** | 59.3, [34.2;108.9] | 16.0, [12.3;20.9] |
| **P2, LS2** | 176.4, [83.9;386.6] | 16.6, [12.3;22.1] |
| **P2, LS3** | 128.3, [57.6;278.7] | 18.7, [14.1;25.9] |

<sup>1</sup> P: parity classes. P1 = first parity, P2 = 2<sup>nd</sup> parity and higher; <sup>2</sup> LS: lactation stage classes: LS1 = days in milk (DIM) 0 to 21, LS2 = DIM 21 to 140, LS3 = DIM 141 to 305; <sup>3</sup> Q: quartiles, Q25 = first quartile, Q75 = last quartile

Moreover, in the first parity, they increased with lactation stage, whereas in higher parities for mid-lactation TD (LS 2, P2), they were highest with median milk losses of 179 kg. When the relative milk losses are considered, the difference between parities and lactation stages almost entirely disappears. Over all parities and lactation stages, the median relative milk losses were 17.2%, with the first and last quartile respectively 12.9 and 23.2%. Similar results are found for each of the P and LS classes, with only milk losses in late lactation higher for both parities (respectively 17.2 and 18.7 %).

### Milk losses at quarter level

#### Milk losses in a fixed window around treatment

From the 3 369 IMI cases with QMY data available, we could unambiguously identify the IQ in 2 190 cases (67.9%). Only these IMI cases were considered for the analysis as they allowed for comparing milk losses between the IQ and the average of the NIQ. From these, the IQ was the left front, right front, left hind and right hind quarter in respectively 558 (25.5%), 559 (25.5%), 520 (23.7%) and 553 (25.3%) IMI cases. This suggests that there is no difference between front or hind quarters in their probability of being infected. For some of the treatment registers, we had information available on the infected quarter as registered by the farmer. The quarter indicated by our data-based method corresponded in 84.0% of those IMI cases with the quarter identified by the farmer.

Table 5 presents the absolute milk losses (in kg) in a fixed window around the TD, separately for IMI cases in which the IQ was a front quarter or a hind quarter. The average absolute milk losses of the NIQ for which IQ was a front quarter includes one front (lower milk yield) and two hind quarters and vice versa. As a result, the median absolute milk losses for front IQ are lower than for hind IQ, and the other way around for the NIQ. Both for front and hind IQ, overall absolute milk losses in the IQ are larger than on average in the NIQ, and this remains true before the TD as well as after the TD, and in all parity and LS classes. Milk yield starts to drop already before the TD in all quarters, but this is more pronounced in the IQ. Furthermore, when analyzing the differences between lactation stages and parities of the IQ, we see similar trends as for the CMY. Milk losses in the IQ are lower in the first lactation stage compared to later stages. There was nearly no difference between the absolute losses in LS2 and 3 in the first parity class, but for IMI cases in higher parities, milk losses in LS2 were clearly higher than those in LS3 for both front and hind IQ. The milk losses in the respective periods before and after the TD are further detailed Table 5.

**Table 5.**
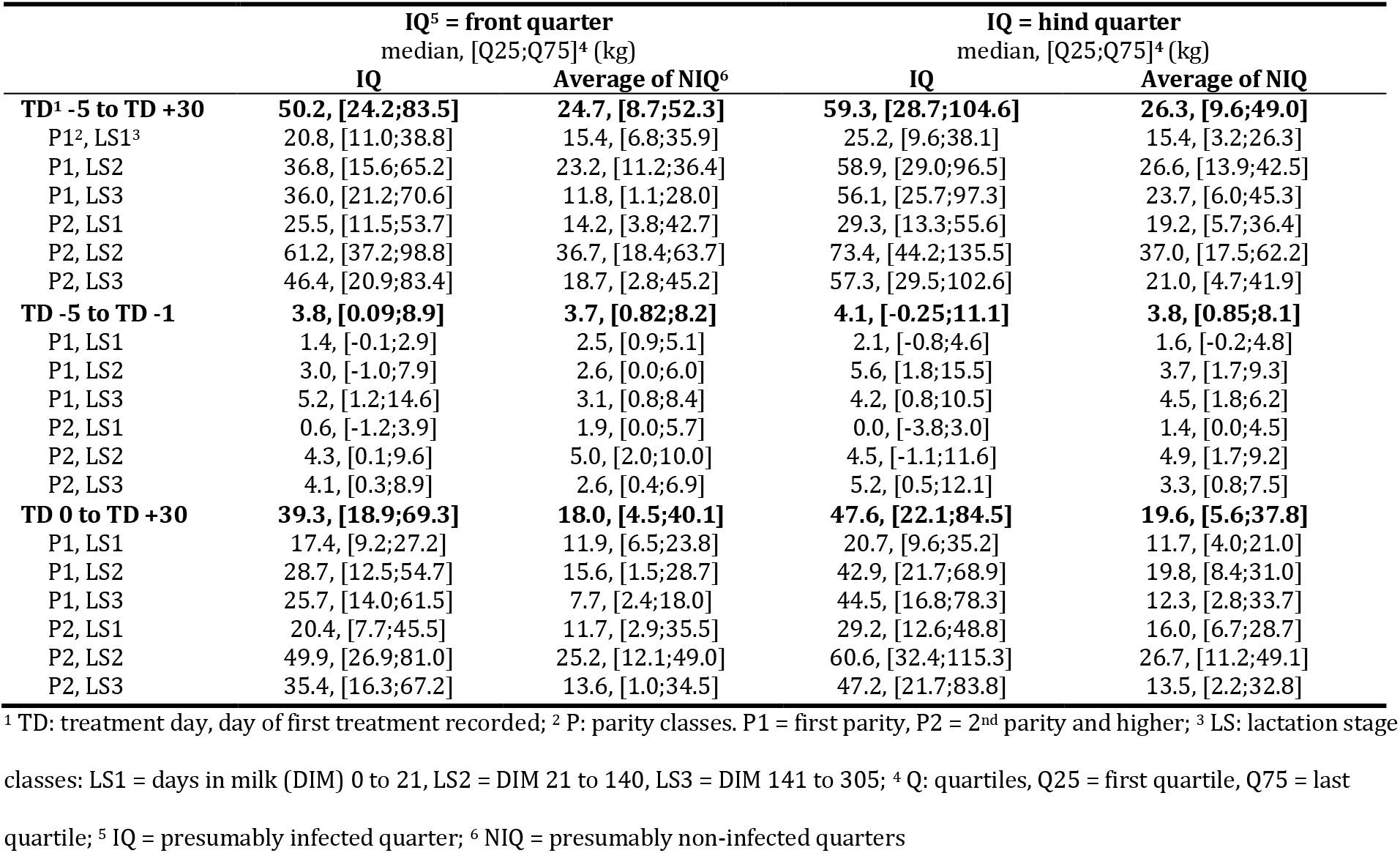
Absolute milk losses at quarter level, for which the presumably infected quarter (IQ) was either a front or a hind quarter, and the average milk losses in their presumably non-infected (NIQ) counterparts in the period of day −5 to 30 relative to the treatment day (TD). The milk losses are expressed as the amount of milk lost, and thus in positive numbers.

|  | IQ<sup>5</sup> = front quarter |  | IQ = hind quarter |  |
| --- | --- | --- | --- | --- |
|  | median, [Q25;Q75]<sup>4</sup> (kg) |  | median, [Q25;Q75]<sup>4</sup> (kg) |  |
|  | **IQ** | **Average of NIQ<sup>6</sup>** | **IQ** | **Average of NIQ** |
| **TD<sup>1</sup> -5 to TD +30** | **50.2, [24.2;83.5]** | **24.7, [8.7;52.3]** | **59.3, [28.7;104.6]** | **26.3, [9.6;49.0]** |
| P1<sup>2</sup>, LS1<sup>3</sup> | 20.8, [11.0;38.8] | 15.4, [6.8;35.9] | 25.2, [9.6;38.1] | 15.4, [3.2;26.3] |
| P1, LS2 | 36.8, [15.6;65.2] | 23.2, [11.2;36.4] | 58.9, [29.0;96.5] | 26.6, [13.9;42.5] |
| P1, LS3 | 36.0, [21.2;70.6] | 11.8, [1.1;28.0] | 56.1, [25.7;97.3] | 23.7, [6.0;45.3] |
| P2, LS1 | 25.5, [11.5;53.7] | 14.2, [3.8;42.7] | 29.3, [13.3;55.6] | 19.2, [5.7;36.4] |
| P2, LS2 | 61.2, [37.2;98.8] | 36.7, [18.4;63.7] | 73.4, [44.2;135.5] | 37.0, [17.5;62.2] |
| P2, LS3 | 46.4, [20.9;83.4] | 18.7, [2.8;45.2] | 57.3, [29.5;102.6] | 21.0, [4.7;41.9] |
| **TD -5 to TD -1** | **3.8, [0.09;8.9]** | **3.7, [0.82;8.2]** | **4.1, [-0.25;11.1]** | **3.8, [0.85;8.1]** |
| P1, LS1 | 1.4, [-0.1;2.9] | 2.5, [0.9;5.1] | 2.1, [-0.8;4.6] | 1.6, [-0.2;4.8] |
| P1, LS2 | 3.0, [-1.0;7.9] | 2.6, [0.0;6.0] | 5.6, [1.8;15.5] | 3.7, [1.7;9.3] |
| P1, LS3 | 5.2, [1.2;14.6] | 3.1, [0.8;8.4] | 4.2, [0.8;10.5] | 4.5, [1.8;6.2] |
| P2, LS1 | 0.6, [-1.2;3.9] | 1.9, [0.0;5.7] | 0.0, [-3.8;3.0] | 1.4, [0.0;4.5] |
| P2, LS2 | 4.3, [0.1;9.6] | 5.0, [2.0;10.0] | 4.5, [-1.1;11.6] | 4.9, [1.7;9.2] |
| P2, LS3 | 4.1, [0.3;8.9] | 2.6, [0.4;6.9] | 5.2, [0.5;12.1] | 3.3, [0.8;7.5] |
| **TD 0 to TD +30** | **39.3, [18.9;69.3]** | **18.0, [4.5;40.1]** | **47.6, [22.1;84.5]** | **19.6, [5.6;37.8]** |
| P1, LS1 | 17.4, [9.2;27.2] | 11.9, [6.5;23.8] | 20.7, [9.6;35.2] | 11.7, [4.0;21.0] |
| P1, LS2 | 28.7, [12.5;54.7] | 15.6, [1.5;28.7] | 42.9, [21.7;68.9] | 19.8, [8.4;31.0] |
| P1, LS3 | 25.7, [14.0;61.5] | 7.7, [2.4;18.0] | 44.5, [16.8;78.3] | 12.3, [2.8;33.7] |
| P2, LS1 | 20.4, [7.7;45.5] | 11.7, [2.9;35.5] | 29.2, [12.6;48.8] | 16.0, [6.7;28.7] |
| P2, LS2 | 49.9, [26.9;81.0] | 25.2, [12.1;49.0] | 60.6, [32.4;115.3] | 26.7, [11.2;49.1] |
| P2, LS3 | 35.4, [16.3;67.2] | 13.6, [1.0;34.5] | 47.2, [21.7;83.8] | 13.5, [2.2;32.8] |

<sup>1</sup> TD: treatment day, day of first treatment recorded; <sup>2</sup> P: parity classes. P1 = first parity, P2 = 2<sup>nd</sup> parity and higher; <sup>3</sup> LS: lactation stage classes: LS1 = days in milk (DIM) 0 to 21, LS2 = DIM 21 to 140, LS3 = DIM 141 to 305; <sup>4</sup> Q: quartiles, Q25 = first quartile, Q75 = last quartile; <sup>5</sup> IQ = presumably infected quarter; <sup>6</sup> NIQ = presumably non-infected quarters

When we express the milk losses relatively in % compared to the expected production, the differences between front and hind IQ disappeared, and therefore were combined them for the rest of the analysis. Figure 4 shows the median and quartiles of relative milk yield dynamics around the TD. We observe that (1) the variability in relative milk losses in the IQ is much larger than in the NIQ; (2) that the median relative losses of IQ on the TD are about 25% higher and (3) that, in contrast to NIQ, full recovery is not reached in most of the IQ within 30 days after the TD. The difference between relative milk losses in IQ and NIQ were significant on day-3 (p = 0.046) and highly significant (P < 0.001) from day-2 to day 30 relative to the TD.

**Figure 4.**
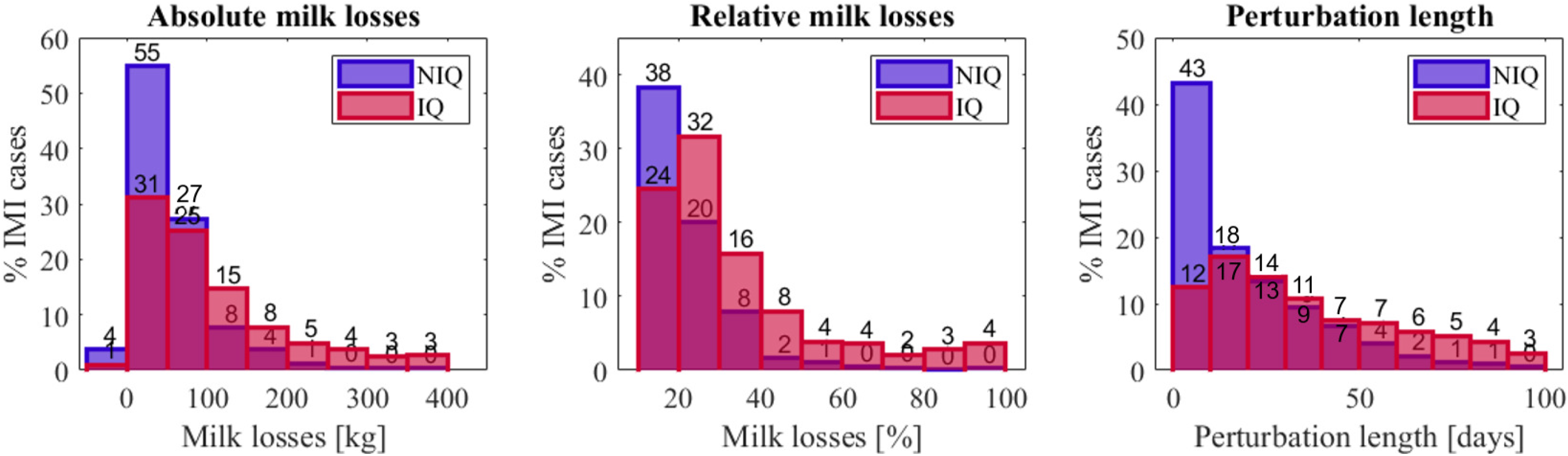
Relative milk losses in the presumably infected (IQ) and the average of the non-infected quarters (NIQ) in the 36 days around treatment. On day −3, the asterisks indicate that the difference between IQ and NIQ is significant with P < 0.05, while on day −2 to 30, the difference is significant with P < 0.001

### Milk losses in perturbations

In 1 878 out of the 2 190 (85.7%) IMI cases for which the IQ was identified, at least one quarter was perturbed at the TD. From these IMI cases, 31% had a perturbation in only one quarter (the IQ), 19% in 2, 20% in 3 and 30% in all 4 quarters. In 35.3% of these IMI cases, the deviation in CMY was too small to meet the perturbation criteria used for that data, explaining the higher proportion of perturbations identified at quarter level compared to the cow level. When a perturbation was found in more than one quarter, the start DIM of these perturbations was for 25% on exactly the same day and in 35% they started within 1 to 5 days from each other. In 18%, the start of the overlapping perturbations was between 5 to 10 days apart and the remaining 22% started more than 10 days apart.

Figure 5 shows the absolute and relative milk losses per perturbation and the perturbation length for the IQ (red) and the NIQ (blue). From this figure and as shown per lactation stage and parity class in Table 6, we observe that the milk losses, both in absolute numbers and relatively, are higher for the IQ than for the NIQ. This confirms the results of the analysis in a fixed window around the TD. In 59% of the IMI cases, the average absolute milk losses in NIQ, independent whether it concerns a hind or front quarter is below 50 kg, whereas this is true for only 32% of the IQ. Expressed as relative losses (middle panel in Figure 5), 38% of the IMI cases have NIQ milk losses less than 20%, whereas most IQ (32%) lose between 20 to 30% compared to the expected production. Also, the average perturbation length of the IQ is higher than in the NIQ, with for the NIQ, 43% of the perturbations shorter than 10 days, where this is only 12% for the IQ.

**Figure 5.**
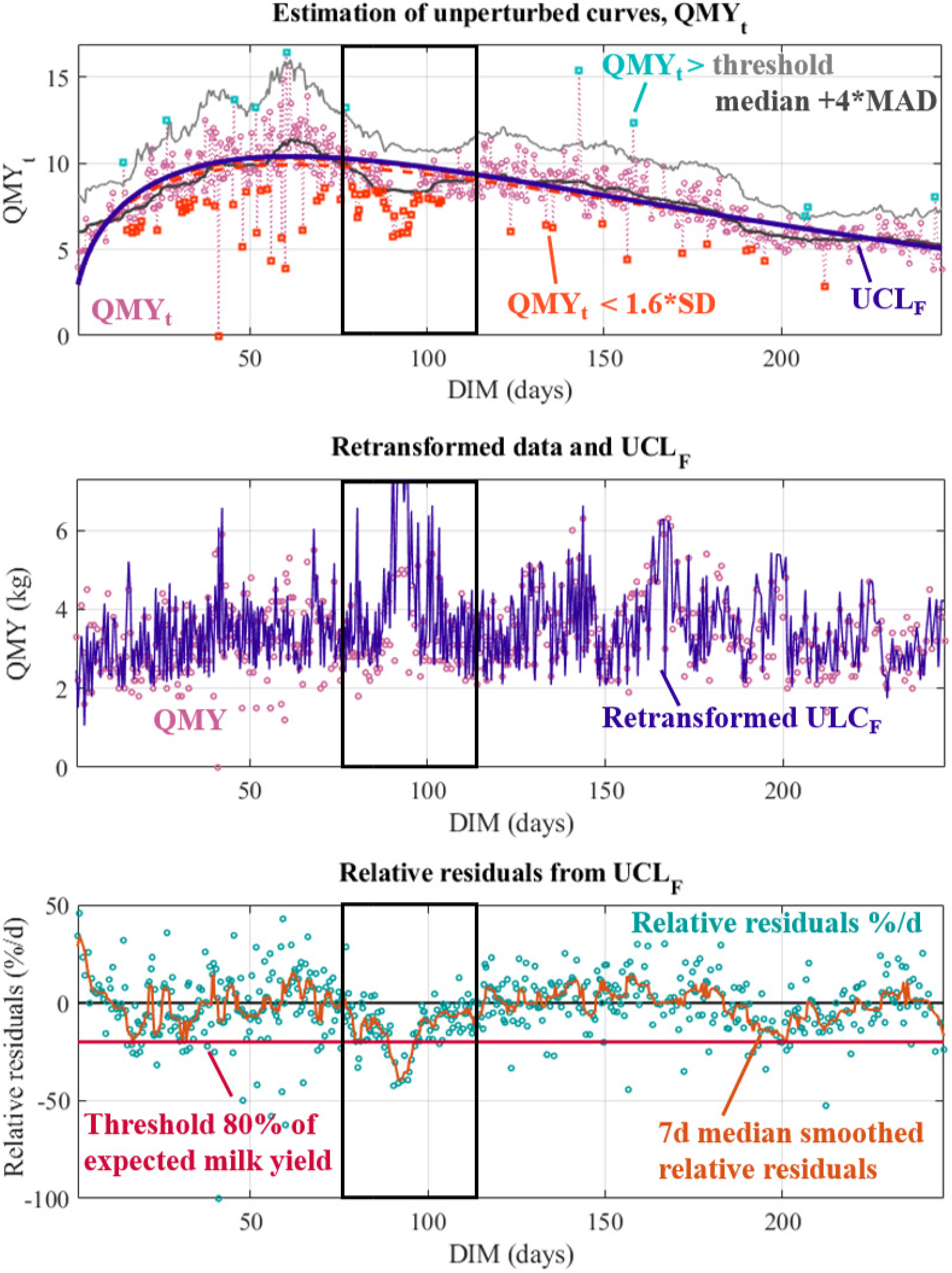
Distributions of milk losses and perturbation length in presumably infected (IQ) and non-infected quarters (NIQ).

**Table 6.** Absolute milk losses in perturbations associated with a IMI treatment in presumably infected (IQ) and non infected (NIQ) quarters, and the corresponding relative losses per day. The milk losses are expressed as the amount of milk lost, and thus in positive numbers.

|  | Absolute milk losses (kg) |  |  |  | Relative milk losses (%) |  |
| --- | --- | --- | --- | --- | --- | --- |
|  | IQ<sup>4</sup> = front quarter<br>median, [Q25;Q75]<sup>3</sup> |  | IQ = hind quarter<br>median, [Q25;Q75] |  | IQ = front and hind<br>median, [Q25;Q75] |  |
|  | IQ | Average of NIQ<sup>5</sup> | IQ | Average of NIQ | IQ | Average of NIQ |
| Overall | 75.7, [35.5;143.3] | 41.8, [19.3;73.5] | 92.7, [45.3;199.9] | 40.9, [20.2;74.7] | 26.0, [19.0;39.3] | 14.7, [8.7;22.0] |
| P1<sup>1</sup>, LS1<sup>2</sup> | 22.0, [11.9;49.0] | 18.8, [8.4;35.8] | 31.6, [17.7;59.8] | 18.7, [12.7;28.3] | 28.0, [20.6;37.2] | 15.1, [10.0;23.0] |
| P1, LS2 | 65.0, [33.2;16.43] | 42.5, [19.6;64.7] | 97.7, [33.5;145.8] | 36.0, [18.6;61.8] | 24.9, [19.2;32.8] | 12.1, [6.1;20.8] |
| P1, LS3 | 60.1, [32.6;114.1] | 22.5, [8.4;45.2] | 88.3, [37.7;173.4] | 34.2, [15.1;60.7] | 30.5, [20.4;49.6] | 13.1, [7.1;21.5] |
| P2, LS1 | 30.5, [17.8;49.5] | 25.7, [11.5;48.1] | 40.7, [19.8;74.0] | 26.2, [13.1;46.5] | 24.5, [18.4;31.8] | 16.5, [11.5;21.4] |
| P2, LS2 | 86.9, [42.1;157.6] | 51.7, [30.1;90.0] | 106.7, [58.2;245.4] | 54.0, [28.0;88.7] | 24.0, [18.4;34.6] | 14.8, [9.6;22.0] |
| P2, LS3 | 78.1, [34.2;151.0] | 39.4, [15.3;67.9] | 101.5, [51.6;211.7] | 36.5, [16.7;66.0] | 28.2, [19.9;44.2] | 14.7, [7.7;22.6] |

<sup>1</sup> P: parity classes. P1 = first parity, P2 = 2<sup>nd</sup> parity and higher; <sup>2</sup> LS: lactation stage classes: LS1 = days in milk (DIM) 0 to 21, LS2 = DIM 21 to 140, LS3 = DIM 141 to 305; <sup>3</sup> Q: quartiles, Q25 = first quartile, Q75 = last quartile; <sup>4</sup> IQ = presumably infected quarter; <sup>5</sup> NIQ = presumably non-infected quarters

Table 6 gives the milk losses of the perturbations at quarter level per lactation stage and parity class. From this table, we see that the median losses in perturbations in the IQ are higher compared to the NIQ for the same class, and that absolute losses in mid lactation (LS2) exceed the losses in early and late lactation cases (LS1 and LS3 respectively). Moreover, when the IQ is a front quarter, the absolute milk losses in perturbations are lower than when the IQ is a hind quarter. The relative milk losses in the IQ, independent of the quarter position, are almost double the relative losses of the NIQ.

## DISCUSSION

### Study design and objectives

This study investigated milk losses associated with treatments of IMI at cow and quarter level on farms with an AMS. Its approach is unique because (1) we used commercial farm data from multiple dairy operations and 2 AMS brands, (2) analyzed naturally occurring IMI and thus did not need to rely on controlled experiments limiting the number of IMI cases for both feasibility and animal welfare issues, (3) investigated both the milk losses at cow and at quarter level for different parities. The latter allowed to focus on differences between more and less severely affected quarters providing unique insights in both local and systemic effects of the infection caused by e.g. decreased feed intake.

We are aware that the use of commercial data and treatment registers involves a risk of including false registrations and erroneous case identification in the analysis (Stevens et al., 2016). Nevertheless, we reasoned that although the chance of missing IMI cases was fairly significant, when a case was registered, the probability that an IMI had truly been treated was very high. The results of this study demonstrate that in most IMI cases the milk production is indeed suppressed around the recorded treatments. Because of the scale of the study and the high number of IMI cases included, false registrations are assumed to minimally impact our results. Unfortunately, no concrete data on false recordings to compare with are available in literature.

This study included data from over 30 commercial farms, nonetheless we could not compare them because of significant differences in quality of the treatment registers and variability in the time period for which milk production data and treatment registers were available. Accordingly, it was not possible to make valid conclusions on the performance of individual farms and how milk production is influenced by IMI. Still, the present joint analysis gives a valuable overview of the impact of IMI on milk losses. A potentially interesting approach for comparing farms could be to start from the quarter milk production data to identify QMY perturbations that are likely caused by IMI. This way, a better idea of the IMI incidences and related milk losses could be obtained including all the data available, and differences in farm performance can be studied without relying on farmers’ recording efforts. This is a promising path for future research.

### Methodology

To study milk losses, an accurate estimation of the expected unperturbed milk yield for a certain lactation is of utmost importance. Previous work showed that mastitis influences the lactation curves, and therefore classical lactation models including the entire time series cannot be used as such (Andersen et al., 2011). Several authors have recently developed methodologies that predict the unperturbed milk yield trajectory of a cow (Adriaens et al., 2020, 2021; Poppe et al., 2020; Ben Abdelkrim et al., 2021), from which the residuals can be used to quantify deviations. The methodology that we developed previously has the advantage of being applicable for QMY trajectories too, which allowed us to use an similar method at both the cow and the quarter level in this study. Data-based estimation of unperturbed trajectories also has disadvantages: when an infection permanently damages the udder and the milk yield never returns to its original potential, it will not be corrected for by the methodology. This can explain the differences in milk losses found here compared to those reported in studies applying a case-control methodology (Heikkilä et al., 2018). Leitner et al. (2020) found altered histology of the mammary gland caused by mastitis, potentially causing a more permanent lower production by the inflamed udder quarters. Analysis of milk yield trajectories across different quarters and over lactations might provide insight into the normal consistency and proportionalities between the QMY, especially considering the high day-to-day variation (Forsbäck et al., 2010). This can help to better understand how often permanent udder damage happens and what the effect is on the true milk production. Penry et al. (2018) showed that there are significant associations between quarters for individual milking events, but they did not consider longitudinal data characteristics. Nevertheless, our earlier work (Adriaens et al., 2018) showed that both the peak yield and the persistency in unperturbed milk yield curves differ across front and hind quarters, and that the variability in quarter lactation shapes suggest that udder asymmetry and past infections may cause the expected milk yield of the left and right udder quarters to be different.

True perturbations were discriminated from normal fluctuations in yield by using pre-set criteria for length and depth of the model residuals, defined as the difference between the actual production and the unperturbed lactation curves. Although these criteria may not capture very subtle perturbations, previous research from Gröhn et al. (2004) and Hertl et al. (2014) showed that IMI typically impact the milk yield well above the applied criteria. Including the perturbations in the analysis has the advantage that a better estimation of the true onset, duration and impact of the infection on the milk yield dynamics can be obtained.

### Milk losses at cow level

In this study we analyzed milk losses associated with an IMI treatment both in a fixed window around the TD and in perturbations including the TD. Our results correspond in general well to previously reported results, but because of unalike milk production data frequencies and the selection procedure of IMI cases, direct comparisons were not possible. Still, similar to other authors, we found a high variability in the milk losses associated with IMI. Part of this variability can be attributed to differences in causal pathogen or clinical severity, which we did not take into account in this study. Additionally, also differences in standard operating procedures of the farmers for registering and treating cows can explain this observation (Heikkilä et al., 2018). Many studies report effects of health status (Hertl et al., 2014), metabolic status (Gross et al., 2020), persistency of the infection (White et al., 2010) and lactation stage (Heikkilä et al., 2018) on the milk losses. In this study, we discriminated between lactation stage and parity classes, but information on the pathogens was not available. Bar et al. (2008) found that both parity and lactation stage indeed alter the effect of IMI on the milk losses, and their estimates were in a similar order of magnitude as in this study, but estimated with a lower granularity of the data. Additionally, Shim et al. (2004) demonstrated that the actual milk losses also depend on the treatments applied, whereas Wilson et al. (2004) found an effect of simultaneous health problems other than mastitis. For many IMI cases, our estimated milk losses were lower than those reported previously in comparison with healthy herd mates, when milk yield measured at a lower frequency was used or when the milk losses were calculated based on an increase in somatic cell count. In the study of Wilson et al. (2004), the milk production losses were estimated around 600 kg for higher parity cows being infected mid-lactation, which is higher than what we found with our analysis, probably because we did not select a certain type of IMI neither looked across different health statuses that might increase the milk losses. Based on their simulations, Van Soest et al. (2016) estimated milk losses for subclinical mastitis and clinical mastitis to be respectively 107 and 306 kg. Given the present variability, these estimates correspond rather well with what we found in this study. Besides the differences in study design and case selection, other differences in milk losses can be explained by the different method by which we quantified the milk losses. In this study, the historical data and expected production dynamics of the treated cows themselves were used, and no comparisons were made with e.g. production levels of the herd or cows with similar parity and lactation stage, neither did we apply fixed percentages or formulas based on e.g. the cell count level. Our methodology might have the disadvantage that milk losses of cases in which damage of the udder tissue causes a permanent milk production drop are probably underestimated.

For monitoring purposes, analysis at cow level might not suffice because of (1) a dilution effect of non-infected quarters on the milk production, not allowing for the separation of local and systemic losses (Boland et al., 2013), and (2) compensatory effects between quarters that have been demonstrated (Hamann and Reichmuth, 1990; Skarbye et al., 2018). A solution to this problem is the analysis of milk losses at quarter level.

### Milk losses at quarter level

Zhao and Lacasse (2008) described how IMI, besides the systemic impact of the disease, also have a local effect on the mammary tissue due to the inflammation reaction and the pathogen’s toxins. This diversification opens opportunities for studying the local and more general effects of the infection, but also for the development of new monitoring tools using quarter level dynamics. With the increased use of AMS, these QMY data become more and more available (Barkema et al., 2015). Indeed, this study demonstrated differences in milk losses between IQ and NIQ, from already days before the TD. Based on the differing number of quarters in which a perturbation was found, it appears that in milder cases, part of the lost milk production in the IQ seems to be compensated in the NIQ, which is not captured with the analysis at cow level. Additionally, the effect of the IMI on the IQ lasted much longer than on the other quarters that only suffered from the systemic impact of the infection. We thereby confirmed the study of Paixão et al. (2017) who argued that also the adjacent quarters’ health status and milk composition is altered by mastitis, although with a lower severity. In the case-control studies conducted by Gonçalves (2018, 2020), the causative pathogen was found to strongly affect the ratio of quarter milk yields, next to the overall milk yield, in cows with subclinical mastitis. For example, they reported daily milk losses between 0.8 and 1.3 kg per quarter (both infected and non-infected), corresponding well to the results of our study.

### Applications and future research

In addition to a very detailed overview of the effect of IMI on milk production at the cow level, this study also clearly reveals the potential of monitoring the production dynamics at quarter level in the context of udder health. Many authors have already shown that mastitis remains one of the most important diseases compromising sustainability of dairy farms, and the economic impact is immense (Hogeveen et al., 2011; van Soest et al., 2016). Milk losses are an important part of the udder health failure costs, and the development of dedicated algorithms for monitoring recovery and cure has a huge potential to increase the relative advantage of AMS systems. These algorithms can for example rely on a change in milk yield dynamics, as demonstrated in this study. Stone (2020) argues that adoption of new PLF innovations indeed depends on this relative advantage, their compatibility with current monitoring systems, their complexity and their observability. Cabrera et al. (2020) reason that the key opportunities for this lie in real-time continuous decision support based on data already available on many modern dairy farms. For example, combining the methodology of the present study with e.g. online modelling and prediction (Adriaens et al., 2018) and a synergistic control method (Huybrechts et al., 2014) could provide the necessary leverage for implementation and adoption of a quarter-based monitoring method maximizing the benefit of already available on-farm sensors. This can aid in changing a farmer’s attitude towards udder health management and antimicrobial use, with an improved sustainability and a better socially accepted dairy production as a result (Deng et al., 2020).

## CONCLUSIONS

This study confirms the high impact of IMI on the milk production of dairy cows in farms with an AMS. Application of the methodology for estimating the unperturbed milk yield allowed to analyze milk losses at the level of both the cow and the udder quarters. Next to the analysis of milk losses in a fixed window around the treatment, also milk losses associated with perturbations including the treatment day were considered. This allowed to estimate and take into account the true duration of the perturbation. The analysis at quarter level showed clear differences between infected and non-infected quarters, revealing the potential for developing novel udder health monitoring systems that make use of commercially available on-farm data.

## ACKNOWLEDGEMENTS

Ines Adriaens received funding from a KU Leuven postdoctoral mandate, grant number PDM/19/132 (Leuven, Belgium). The data from this study were collected in the format of farm software back-up files by the authors, and the resulting database is owned by the authors. We thank dr. DVM. Sofie Piepers (Ghent University, Department of Reproduction, Obstetrics and Herd Health, M-team and Mastitis and Milk Quality Research Unit, Belgium) for her help in the data collection during the farm visits. Furthermore, this study is funded by VLAIO as an LA-trajectory (Brussels, Belgium), grant number HBC.2016.0774. We thank Carmen Adriaens (Harvard Medical School and Massachusetts General Hospital, Boston, USA) for her critical reading of the manuscript. The authors have not stated any conflicts of interest.

